# The neural bases of resilient cognitive systems: Evidence of variable neuro-displacement in the semantic system

**DOI:** 10.1101/716266

**Authors:** JeYoung Jung, Grace E. Rice, Matthew A. Lambon Ralph

## Abstract

The purpose of this study was to initiate exploration of an equally-important research goal: what are the neurocomputational mechanisms that make these cognitive systems “well engineered” and thus resilient across a range of performance demands and to mild levels of perturbation or even damage? We achieved this aim by investigating the neural dynamics of the semantic network with two task difficulty manipulations. We found that intrinsic resilience-related mechanisms were observed in both the domain-specific semantic representational system and the parallel executive control networks. Functional connectivity between these regions was also increased and these increases were related to better semantic task performance. Our results suggest that higher cognitive functions are made resilient by flexible, dynamic changes (variable neuro-displacement) across both domain-specific and multi-demand networks. Our findings provide strong evidence that the compensatory functional alterations in the impaired brain might reflect intrinsic mechanisms of a well-engineered neural system.

## Introduction

One critical feature of any well-engineered system is its resilience to variable performance demands, perturbation and minor damage. How resilience is achieved in higher cognitive systems is important both for cognitive and clinical neuroscience. Although rarely considered in laboratory-based explorations of higher cognition (beyond executive function where demand variations are inherently important), in everyday life we are faced with and are resilient to variations in task difficulty, degraded stimuli, etc. Likewise, after partial brain damage or perturbation, participants can sometimes show impressive resilience and recovery. The current study investigated the mechanisms that support resilience in the domain of semantic cognition.

Semantic cognition allows us to use, manipulate and generalize knowledge. This is a crucial function for communication (verbal and nonverbal) and activities of daily living (e.g., object use) (1). It can be decomposed into two components: *semantic representation* – the long-term representation of concepts/semantic memory and *semantic control* – mechanisms to generate time- and context-appropriate semantic behaviours (2). The neural basis of semantic cognition reflects a large-scale network across the frontal, temporal and parietal cortex (3, 4). Convergent evidence suggests that the ventrolateral anterior temporal lobe (ATL) is the centre-point of a transmodal hub that supports semantic representation (5-9), whereas prefrontal and temporoparietal cortices are involved in controlled retrieval of semantic knowledge (10-13).

There are two existing sources of evidence for resilience in the semantic system. The first comes from patients with unilateral ATL resection/damage (either left or right) who exhibit mild semantic impairments (reflected in slower response times or reduced accuracy on demanding semantic tasks) but perform much better overall than patients with bilateral ATL resection/damage (14-16). The same pattern was shown in the seminal investigations of unilateral versus bilateral ATL resections in non-human primates (17, 18) and one human case (19), in which unilateral resection resulted in transient multimodal associative agnosia, whereas bilateral resection caused severe, chronic deficits. Whilst these studies show that the semantic system is somewhat robust against partial damage (14, 20), it remains unclear what neural mechanisms underlie this resilience.

The second line of evidence comes from recent studies that combined fMRI and transcranial magnetic stimulation (21, 22). Using a “perturb-and-measure” approach (23), inhibitory repetitive TMS (rTMS) was delivered over the left ATL and the resultant behavioural and neural changes were measured. rTMS over the left ATL decreased regional activity in the target site yet increased activity at the contralateral ATL. This upregulation in the right ATL contributed to residual semantic performance (stronger activity was associated with faster reaction times). Furthermore, an effective connectivity analysis revealed that, after the left ATL stimulation, there was increased connectivity from the right ATL to the left ATL (22).

As well as resultant changes within the ATL-based representational system, functional neuroimaging and patients studies indicate that there is also potentially important upregulation of the semantic control regions (24, 25). Furthermore the distributed network for semantic control, itself, also seems to exhibit dynamic, resilient-related changes in that rTMS to IFG generates compensatory increased activation in the strongly-connected pMTG (26-28). These studies suggest that the robustness of semantic system can be attributed to the involvement of semantic control regions.

The limited data on this topic to date principally centre on patient or TMS investigations – i.e., examining the brain after damage or perturbation. The central aim of this study was to test whether these dynamic changes are specific to the impaired brain (indicating compensatory changes that are triggered by brain damage/perturbation) or whether they reflect intrinsic mechanisms of a well-engineered cognitive system. A core tenet in engineering is to design a system that is resilient to functional stresses, as well as to balance performance and energy costs. It seems likely that neurocognitive systems are also designed: a) to be tolerant to variable levels of performance demand; and b) to titrate performance against metabolic energy demands. Accordingly, under standard levels of performance demand the full semantic system will be down-regulated to save energy but have spare capacity that can be utilised when the situation necessitates it (29) (known as ‘variable displacement’ in modern designs of combustion engines) (30). We tested this possibility by investigating the neural dynamics of semantic system in healthy participants at different levels of performance challenge. To ensure generalisation across different types of challenge, we manipulated task difficulty in two different ways (stimulus difficulty vs. response timing). In the response timing experiment, we also compared the results for verbal (written words) and nonverbal (picture) semantic processing. If the changes observed in previous studies of patients or rTMS reflect intrinsic features of a well-engineered system then we expected to see the same types of dynamic changes in the healthy system under performance pressure. These would include changes in the ATL semantic representational system and the prefrontal-pMTG semantic and multi-demand executive control networks.

## Materials and Methods

### Participants

Forty-one healthy participants were recruited for the study. Twenty-one participants (7 females, mean age, 22 ± 3.1 years) participated in the stimulus manipulation (SM) experiment and twenty participants in the response timing manipulation (RM) experiment. All participants were right-handed, native English speakers who had normal or corrected-to-normal vision. They received a detailed explanation of the study and gave written informed consent prior to the experiment. The experiment was approved by the local ethics committee.

### Experimental design

Participants underwent two tasks with two different level of difficulty (easy vs. hard): a semantic task and a matched control task. We manipulated task difficulty in two different ways: stimulus manipulation (SM) and response time manipulation (RM) (Fig.1). The RM data have been reported in a recent examination of the semantic network in patients with ATL resection (31).

**Figure 1.**
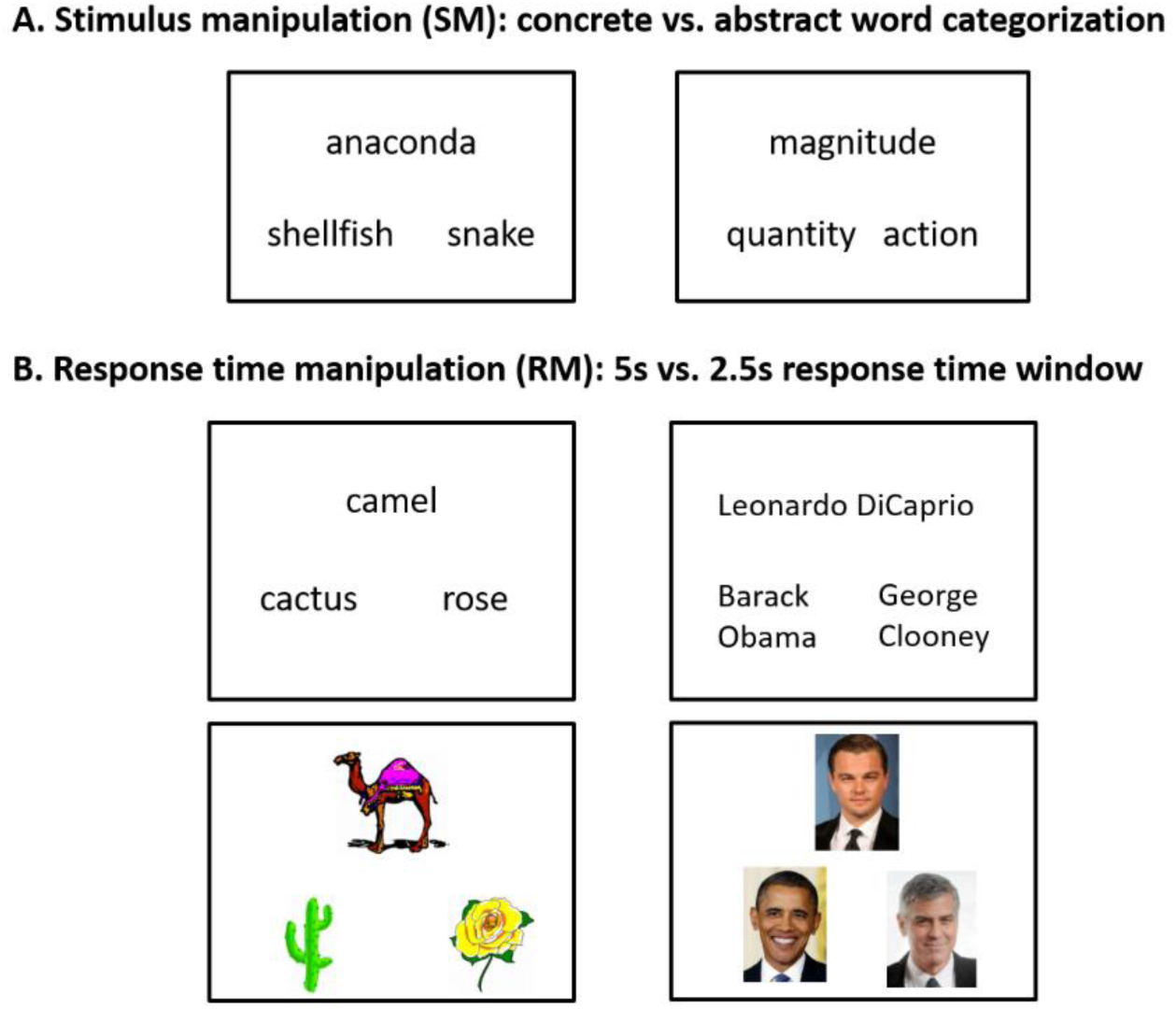
Experimental design. A) Stimulus manipulation (SM). B) Response time manipulation (RM)

In the SM experiment, participants performed a category judgement task. The stimuli for the easy condition were from the *Levels of Familiarity, Typicality, and Specificity (LOFTS)* semantic battery (32). The concrete words covered a variety of categories, including animals, vehicles, tools, foods, and plants. The stimuli for the hard condition were selected from a previous study rating abstract nouns according to conceptual features (i.e., emotion, sensation, action, thought, time, quantity, morality, social interaction, and teaching) (33). 335 abstract nouns were selected and six native English speakers rated them for their category. In this study, we used abstract nouns that all six raters agreed to a category. Participants were asked to indicate which of two categories was appropriate for a target. In each trial, three words were presented on the screen, a target on the top and two choices at the bottom (Fig.1A). A pattern matching task was used as a control task. The items for the control task were generated by visually scrambling items from the semantic task. Each pattern was created by scrambling each item into 120 pieces and re-arranging them in a random order. Participants were asked to select which of two patterns was identical to a target pattern. In the hard condition, the patterns for a right answer were 180° rotated.

In the RM experiment, participants performed the Camel and Cactus test (CCT) (34) and an occupation matching task for famous people as semantic tasks (31) (Fig. 1B). On each trial, three items were presented on the screen, a probe concept at the top and two choices at the bottom. Participants were asked to decide which of the two alternatives was semantically related to the target for the CCT and to indicate which of two alternatives had the same occupation for the people matching task. This resulted in four semantic conditions (CCT [word], CCT [picture], Famous Faces, Famous Names). In the occupation matching task, all items shared the same gender. To investigate the effects of the modality, both tasks were presented either as written words and pictures. The same pattern matching task was used as a control task. In order to manipulate the difficulty in response, the stimuli were presented with 2.5 s response window for the hard condition and 5 s for the easy condition.

### Experimental procedure

The total scan time of the SM experiment was about 10 minutes. During scanning, stimuli were presented in a block design and each block contained 4 trials from one experimental condition (semantic: concrete and abstract words and control: patterns and rotated patterns). Each stimulus and the response screen were presented for 5000ms, with an inter-stimulus interval of 500ms. The four experimental conditions were sampled six times in a counterbalanced order, giving a total of 24 blocks.

The total scan time of the RM experiment was about 8.45 minutes for the easy condition and 4.2 minutes for the hard condition. During scanning, stimuli were presented in a block design and each block contained three trials from one experimental condition. Participants performed eight functional scans containing stimuli from one semantic condition (CCT [word], CCT [picture], famous names or famous faces) and from the matched control condition (scrambled pictures or scrambled words). For the easy functional scans (four scans), each stimulus and the response screen were presented for 5000ms, with an inter-stimulus interval of 500ms and for the hard scans, difficulty was increased by presenting stimuli twice as quickly, at 2500ms intervals. The functional scans were interleaved to avoid any habituation to the speed of presentation.

All participants underwent practice trials before beginning the scan to familiarize them with the tasks. E-prime software (Psychology Software Tools Inc., Pittsburgh, PA, USA) was used to display stimuli and to record responses.

### fMRI data acquisition and analysis

Imaging was performed on a 3T Philips Achieva scanner using a 32-channel head coil with a SENSE factor 2.5. To improve signal-to-noise (SNR) in the ATL, we utilised a dual-echo fMRI protocol developed by Halai et al (35). The fMRI sequence included 42 slices, 96 × 96 matrix, 240 × 240 × 126mm FOV, in-plane resolution 2.5 × 2.5, slice thickness 3mm, TR = 2.8s, TE = 12ms and 35ms. All images were acquired using a tilt, up to 45°off the AC-PC line, to reduce ghosting artefacts in the temporal lobes. In the SM experiment, 190 dynamic scans were acquired and, in the RS experiment, 177 dynamic scans for the easy condition and 88 dynamic scans for the hard condition (all functional scans included two dummy scans, which were excluded). The structural image was acquired using a 3D MPRAGE pulse sequence with 200 slices, in planed resolution 0.94 × 0.94, slice thinkness 0.9mm, TR = 8.4ms, and TE = 3.9ms.

Analysis was carried out using SPM8 (Wellcome Department of Imaging Neuroscience, London; www.fil.ion.ucl.ac.uk/spm). The dual gradient echo images were extracted and averaged using in-house MATLAB code developed by Halai et al (35). Functional images were realigned correcting for motion artefacts and different signal acquisition times by shifting the signal measured in each slice relative to the acquisition of the middle slice prior to combining the short and long echo images. The mean functional EPI image was co-registered to the individual T1-weighted image and segmented using the DARTEL (diffeomorphic anatomical registration through an exponentiated lie algebra) toolbox (36). Then, normalization was performed using DARTEL to warp and reslice images into MNI space and smoothing was applied with an 8mm full-width half-maximum Gaussian filter.

At the individual subject level, contrasts of interest were modelled using a box-car function convolved with the canonical hemodynamic response function. For the SM experiment, four separate regressors were modelled according to task and difficulty (semantic: concrete and abstract words and control: patterns and rotated patterns). For the RM experiment, two separate regressors were modelled in each functional scan: 1) semantic condition (either: CCT [picture], CCT [word], Famous Faces, Famous Names) and 2) control condition (either scrambled pictures or scrambled words).

At the group level, a two-factorial ANOVA with task (semantic vs. control) and difficulty (easy vs. hard) was conducted for the main effect of task and interaction between task and difficulty and T-contrasts were established for the contrast of semantic > control and hard > easy across the experiments. Same analyses were performed according to the modality (words and pictures) and difficulty manipulation (SM and RM). Whole-brain maps were thresholded at p < 0.001 at the voxel level, with a FWE-corrected cluster threshold of p < 0.05, ks > 100.

### Region of Interest (ROI) analysis

An a priori ROI analysis was employed to assess the level of activation in semantic regions including the vATL, IFG, and pMTG. Peak coordinates were taken from previous studies investigating the semantic system: vATL [MNI: −36 −15 −30; 36 −15 −30] (5), IFG (BA 45: p. Tri) [MNI: −45 19 18; 47 23 26] (10), IFG (BA47: p. Orb) [MNI: −45 27 −15; 45 27 −15] (37), and pMTG [MNI: −66 −42 3; 66 −42 3] (5). The right hemisphere ROIs were created using the homologous coordinates. Each ROI was created as a sphere with 8mm radius.

### Functional connectivity (FC) analysis

To investigate changes in vATL functional connectivity to the rest of the brain, we conducted psychophysiological interaction analysis (PPI). The vATL has been reported as the centre-point of a representational hub, which interacts with other semantic control regions and spokes to generate coherent semantic knowledge (1, 6, 38). The PPI analysis describes neural responses in one brain area in terms of the interaction between influences of other brain regions and a cognitive process (39). The PPI analysis employed a design matrix with three regressors: (i) the “psychological variable” representing the cognitive process of interest; (ii) the “physiological variable” representing the neural response in the seed region and (iii) the interaction term of (i) and (ii). To quantify the physiological variable, we extracted the individual time series from the left vATL (lvATL) as a seed region. For the lvATL seed, the time courses were de-convolved based on the model for the canonical hemodynamic response to construct a time series of neural activity, which was the physiological factor. Interaction terms were calculated separately for each experimental condition (semantic: easy vs. hard), as the product between the vector of the condition and the physiological factor. The PPI terms were also been convolved with the hemodynamic response function. Then, we revealed how the hard semantic condition induced functional connectivity change to the corresponding seed region compared with the easy condition. As the PPI is very stringent (33), we used the significance threshold at p < 0.005 at the voxel level, with a FWE-corrected cluster threshold of p < 0.05, ks > 30.

In order to investigate the difficulty effects on the semantic processing between semantic regions, we employed the Functional Connectivity (CONN) Toolbox (http://web.mit.edu/swg/software.htm). This method enables examination of network interactions during each condition of an fMRI task and thus comparison of FC between different conditions (semantic: easy vs. hard). Pre-processed images were entered to the toolbox. Data were filtered using a band pass filter (0.01 < f < 2) to decrease the effect of low-frequency drift. White matter, cerebrospinal fluid, and physiological noise source reduction were taken as confounds, following the implemented CompCor strategy (40). Head motion was taken into account and rotational and translational motion parameters and their first-order temporal derivatives were regressed out. The onset and duration of each experimental condition was supplied to the toolbox so as to extract the connectivity generated for easy and hard semantic processing. The key regions of the semantic network were included in this analysis (vATL, IFG [p.Orb], IFG [p. Tri], and pMTG) and bivariate correlations were calculated between each pair of ROIs as reflections of connections according to the experimental conditions. Planned paired t-tests were performed on the FC between ROIs for the easy and hard semantic conditions (p < 0.05).

## Results

### Behavioural results

A repeated-measures ANOVA with task (semantic vs. control) and difficulty (easy vs. hard) was conducted for each experimental dataset (SM and RM) and the combined dataset (SM+RM). In accuracy, the SM dataset revealed a significant main effect difficulty (F_1, 20_ = 35.10, p < 0.001) and an interaction (F_1, 20_ = 10.09, p < 0.01). The RM dataset showed a significant main effect of difficulty (F_1, 19_ = 42.46, p < 0.001). In reaction time (RT), the SM dataset showed a significant main effect of task (F_1, 20_ = 42.87, p < 0.001), difficulty (F_1, 20_ = 212.94, p < 0.001) and an interaction (F_1, 20_ = 22.04, p < 0.001). The RM dataset showed a significant main effect of difficulty (F_1, 19_ = 199.12, p < 0.001). *Post hoc* paired t-tests revealed that the difficulty manipulation was successful for both datasets and for both tasks (Fig. 2). In the SM dataset, the accuracy was significantly reduced and the RT was increased for the hard condition compared to the easy condition for both semantic and control tasks (p < 0.001). In the RM dataset, the accuracy was significantly decreased in the hard condition relative to the easy condition, whereas the RT was faster in the hard condition due to the response time manipulation (p < 0.001). The combined dataset also showed a significant main effect of difficulty (F_1, 40_ = 78.63, p < 0.001) and an interaction (F_1, 40_ = 9.06, p < 0.01) in accuracy. *Post hoc* paired t-tests revealed that accuracy was significantly decreased in the hard condition for both tasks (p < 0.001). In RT, there was a significant main effect of task (F_1, 40_ = 24.28, p < 0.001) and an interaction between task and difficulty (F_1, 40_ = 7.78, p < 0.001). The results of modality (word and picture) are summarised in Fig. S1.

**Figure 2.**
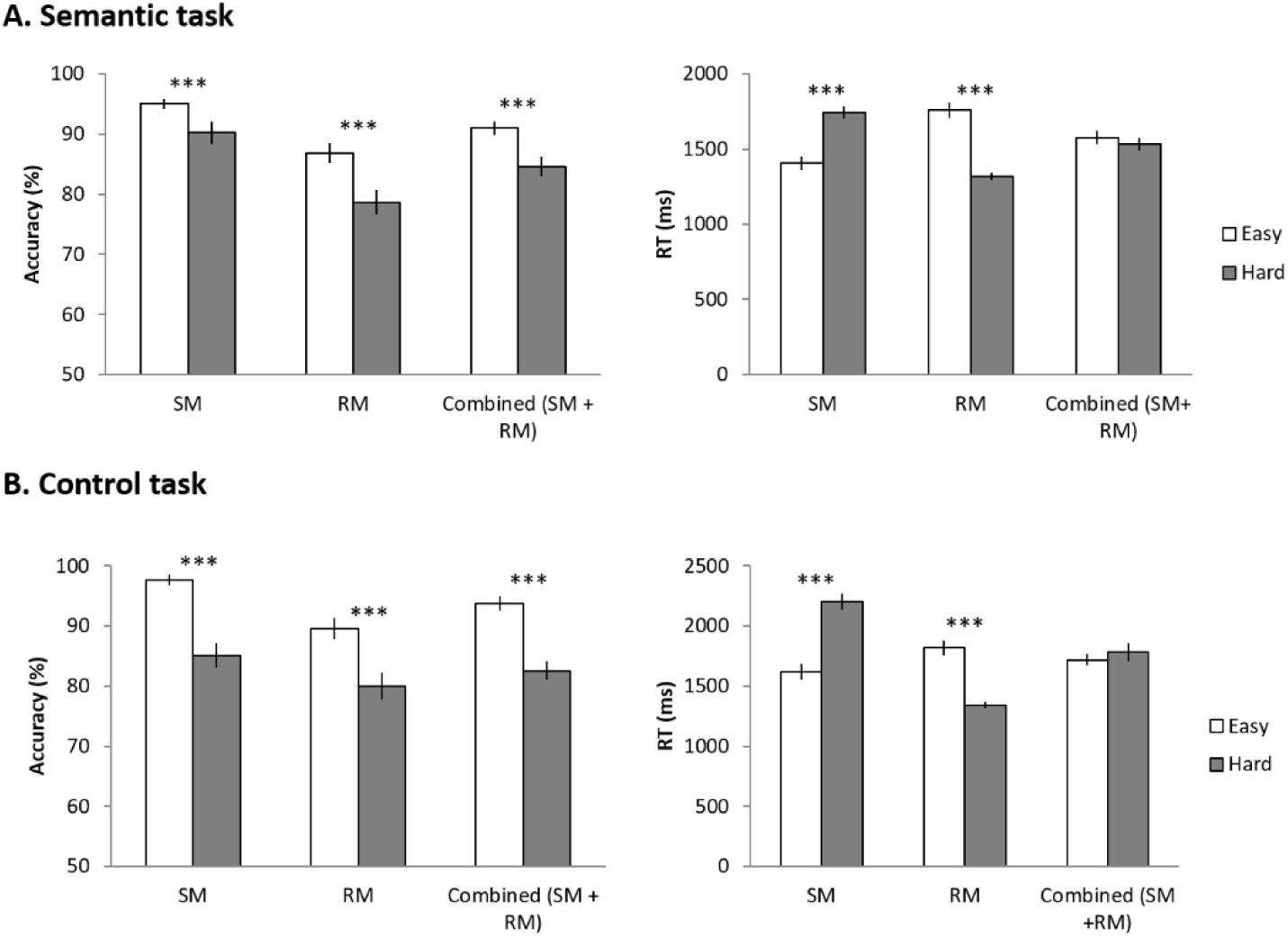
Behavioural results. A) Behavioural results of semantic tasks. B) Behavioural results of control tasks. White bars represent the easy condition performance and grey bras the hard condition performance. Error bars indicate standard errors. ***p < 0.001

### fMRI results

The whole brain analyses revealed that the semantic task (SM+RM) evoked significant activation in the left IFG, vATL, pMTG, fusiform gyrus, medial prefrontal cortex (mPFC), superior frontal gyrus (SFG), and precuneus (PCC) regardless of task difficulty (semantic > control) (Fig. 3A top). The hard semantic condition induced more widespread activation in the same regions and additional activation in the left angular gyrus (AG) and the right ATL (lateral and ventral), anterior MTG (aMTG), and pMTG. It should be noted that the additional activation evoked by the hard condition was not distinct from the pattern of activity during the easy semantic condition if a less stringent threshold was applied (p _unc_ < 0.005 at a voxel level). Even in the easy semantic condition, we found widespread activation in the bilateral ATL, IFG, pMTG, and AG as well as the left precentral gyrus, hippocampus, and basal ganglia (Fig. 3A bottom). The control task (visuospatial processing) activated the bilateral intraparietal sulcus (IPS), inferior parietal lobe (IPL), middle frontal gyrus (MFG), SFG, superior occipital gyrus (SOG), middle occipital gyrus (MOG) and cuneus during the easy condition. The hard control processing was involved in the same regions as well as the right superior parietal lobe (SPL), IFG (p. Opercularis), inferior temporal gyrus (ITG), which overlaps with the easy condition activation map with a lower threshold (Fig. S2). Subsequent ROI analyses demonstrated that more demanding semantic processing significantly increased regional activity in the left semantic regions (IFG [p. orb and p. tri], pMTG, and vATL) compared to the easy semantic condition. Only the right vATL showed a significant up-regulation during the hard semantic condition (Fig. 3B). The same analyses were performed in each dataset (SM and RM), split according to modality, and similar results were obtained (Fig. S3 & S4). The comparison between SM and RM is summarised in Supplementary results and Fig. S5.

**Figure 3.**
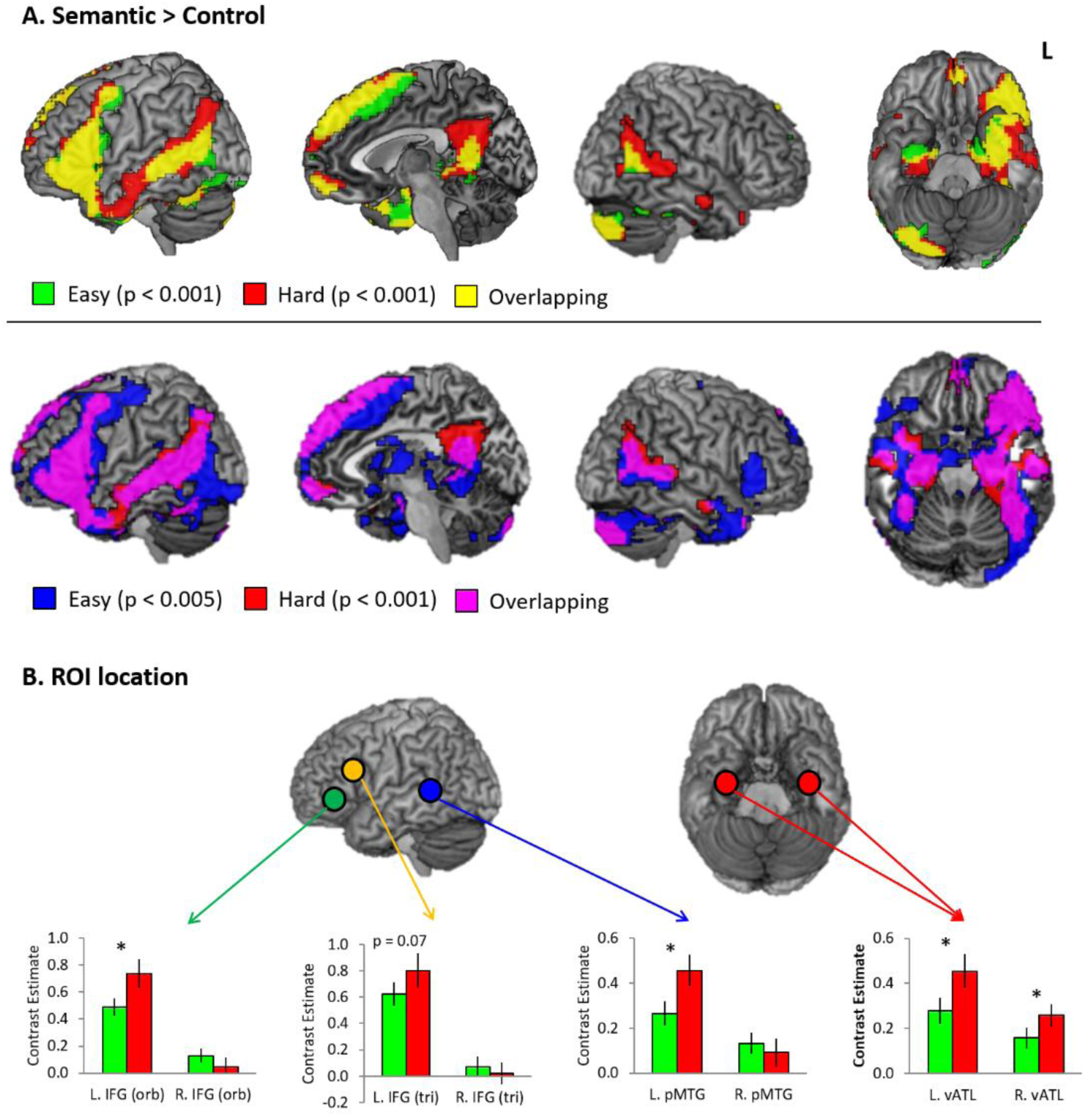
fMRI results. A) Brain activation map of the contrast (Semantic > Control). Green colour indicates the results of the easy semantic condition. Red colour indicates the results of the hard semantic condition. Yellow colour represents overlapping between the easy and hard condition. Blue indicates the result of the easy semantic condition with a lower threshold (p < 0.005). Pink represent the overlapping between the hard condition and easy condition with a lower threshold. B) ROI results. Green bars represent the easy condition and red bars the hard condition. Error bars indicate standard errors. * p < 0.05

Several regions showed an interaction between task and difficulty, many of which overlapped with the areas exhibiting the effect of task and/or difficulty (Fig. 4). Specifically, the left IFG and pMTG, as semantic control regions, showed positive activation for the semantic task and greater activation in the hard semantic condition. The interaction in the AG arose from differential deactivation particularly for the hard visuo-spatial processing. As key regions of the default mode network (DMN), the mPFC and precuneus revealed deactivation for both tasks but a differential difficulty effect was found in these regions according to tasks. The mPFC showed decreased deactivation for hard semantic processing and increased deactivation for demanding visuospatial processing. The precuneus showed the difficulty effect only for visuospatial processing – more deactivation during the hard control condition. The IPS, a key region of the multiple-demanding system (MD) exhibited a task-general effect of difficulty – more activation for demanding condition regardless of tasks. The task-opposite pattern was found in the left lateral occipital cortex (LOC), right ITG and right frontoparietal regions, such that the difficulty effect was observed only for visuospatial processing. These same analyses were repeated on each dataset, split according to modality and the results are summarised in Supplementary results and Fig. S6-7.

**Figure 4.**
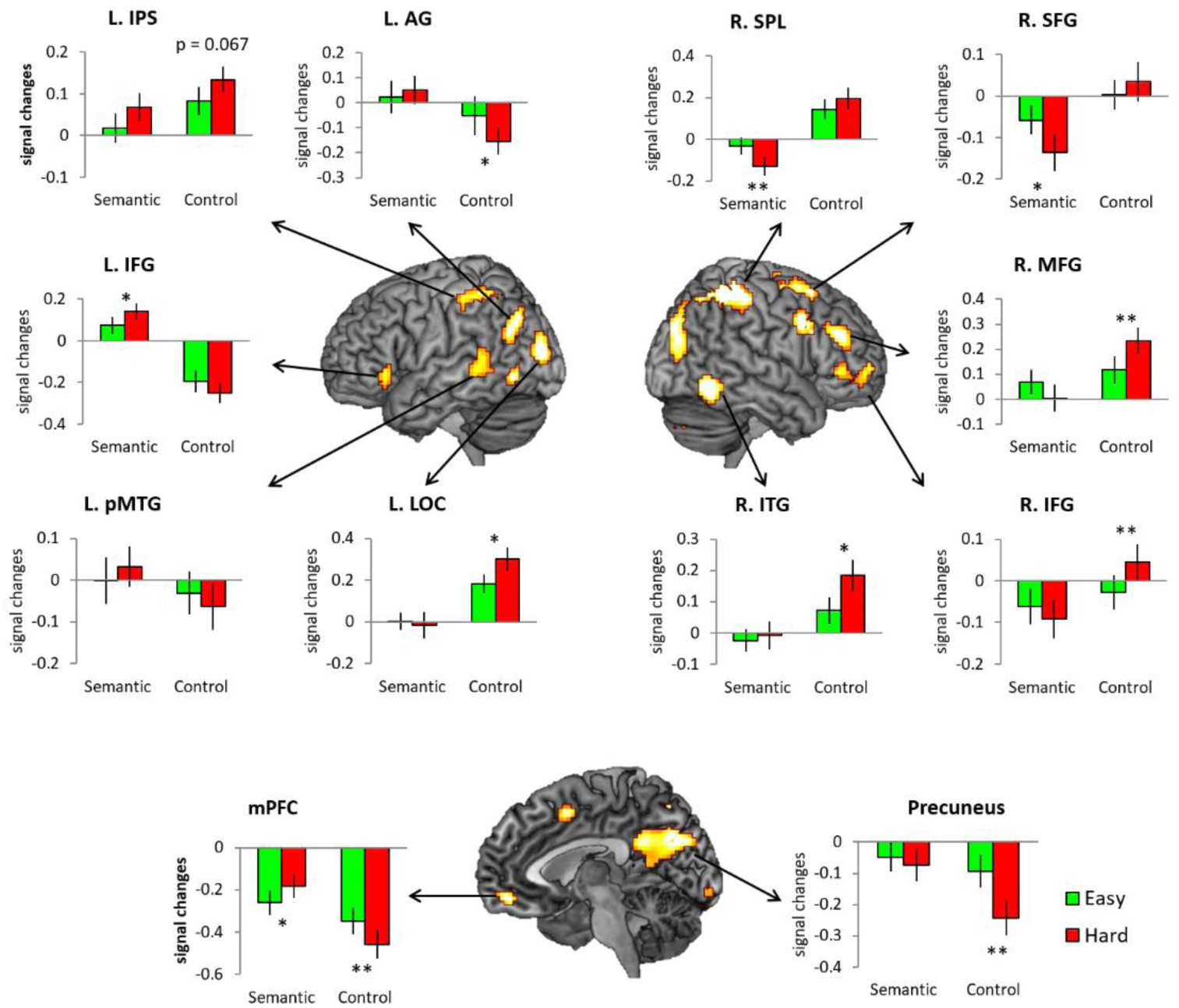
The interaction between task and difficulty. Green bars represent the easy condition and red bars the hard condition. Error bars indicate standard errors. * p < 0.05

### FC results

PPI analyses were conducted to examine how vATL connectivity was modulated by difficulty. The easy semantic condition showed that the left vATL is significantly connected with the bilateral IFG, left orbitofrontal cortex (OFC), supplementary motor area (SMA), visual cortex, and cerebellum. The hard semantic condition increased the left vATL connectivity to the same regions found in the easy condition as well as additional areas including the right vATL, left IPS, MFG mPFC, and insular (Fig. 5A).

**Figure 5.**
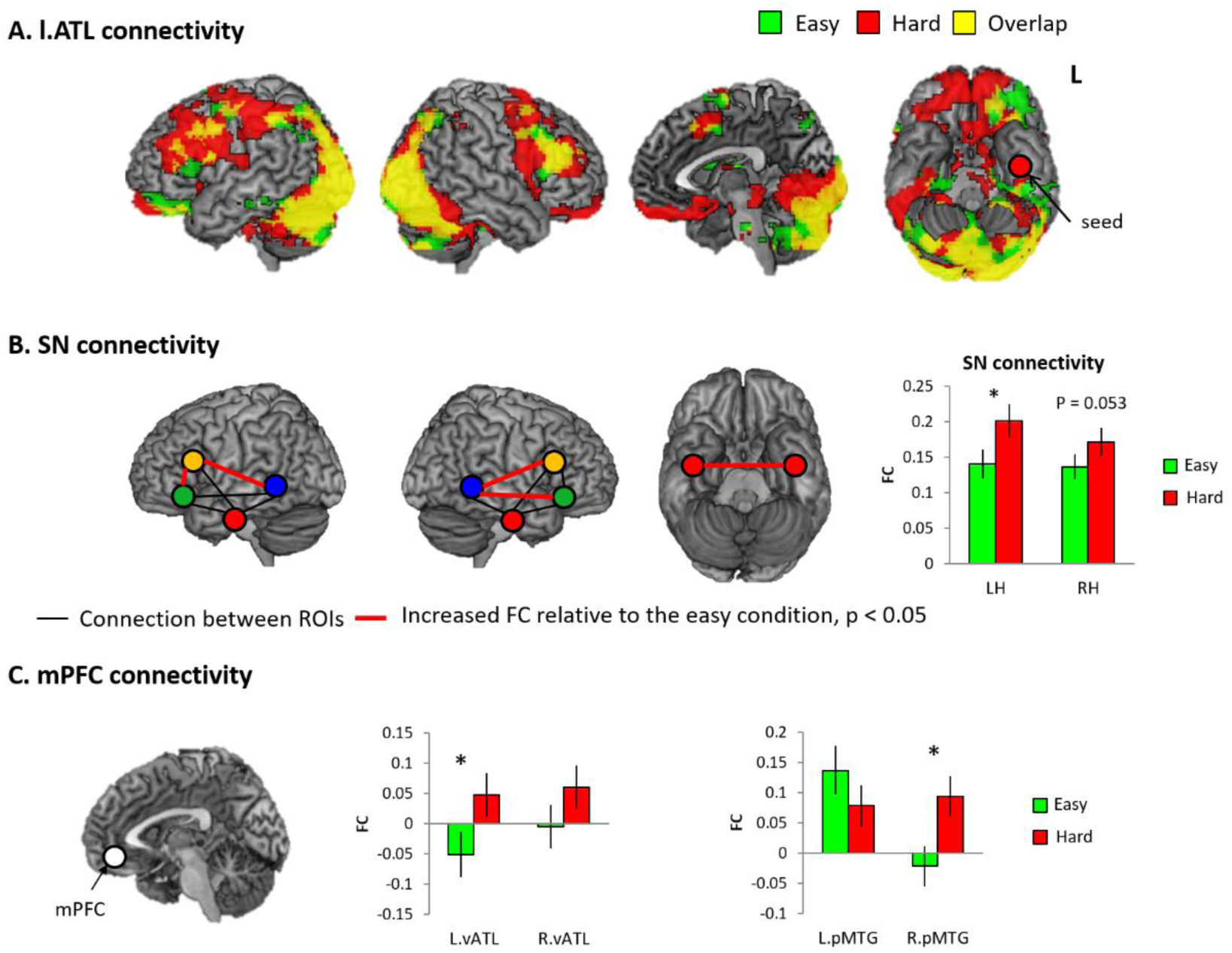
Functional connectivity (FC) analysis. A) The results of PPI with the l.vATL seed. Green colour indicates the results of the easy semantic condition. Red colour indicates the results of the hard semantic condition. Yellow colour represents overlapping between them. B) The results of ROI-to-ROI analysis in the semantic network (SN). Black line represents a connection between ROIs and red line indicates a significantly increased connection between ROIs in the hard semantic condition compared to the easy semantic condition. C) The results of the mPFC connectivity to semantic regions. Green bar indicates the easy semantic condition. Red bar indicates the hard semantic condition. * p < 0.05

We examined how ROI-to-ROI connectivity in the semantic network (SN: IFG [p. orb, p. tri], pMTG, and vATL) changed across the semantic difficulty levels. The FC between the IFG and pMTG in both hemispheres was significantly increased in the hard condition (Fig. 5B). Also, the averaged FC in all semantic regions (SN connectivity) in the both hemispheres was significantly increased in the hard semantic condition.

ROI and PPI analyses revealed there was greater activation in the mPFC during more demanding semantic processing. We explored the semantic difficulty effect in the FC between the mPFC and semantic regions. A 2 × 2 ANOVA with difficulty (easy vs. hard) and hemisphere (left vs. right) was conducted. The results demonstrated that the hard semantic condition significantly increased the mPFC– vATL connectivity (a main effect of difficulty: F_1, 40_ = 4.40, p < 0.05) and the mPFC– right pMTG connectivity (a main effect of hemisphere: F_1, 40_ = 6.41, p < 0.05, interaction: F_1, 40_ = 4.07, p < 0.05) (Fig. 5C). *Post hoc* paired t-tests confirmed the findings. The same analyses were performed in each dataset, split according to modality, and the results are summarised in Fig. S8.

### FC-behavioural correlations

We correlated semantic performance with FC between semantic regions to examine which functional connectivity aligns with the changing semantic demand. Accuracy during the easy semantic condition was positively correlated with the left– right vATL and left vATL–left IFG [p. Tri] connectivity (Fig. 6A). RT in the easy semantic condition was significantly correlated with the left–right vATL, left vATL–left IFG, and left SN connectivity (p _FDR-corrected_ < 0.05) (Fig. 6A). In the hard semantic condition, there were significant correlations between accuracy and the interhemispheric vATL connectivity (left–right vATL) and between accuracy and the vATL–IFG [p. orb, p. tri] connectivity in both hemispheres (p _FDR-corrected_ < 0.05) (Fig. 6B). We did not run the analysis for the RT in the hard semantic condition, as there was no difficulty effect in the combined dataset (SM + RM). Overall, individuals with stronger FC between the semantic regions performed the semantic task better (i.e., faster RT and/or higher accuracy). The same analyses were performed in each dataset, split according to modality, and the results are summarised in Fig. S9.

**Figure 6.**
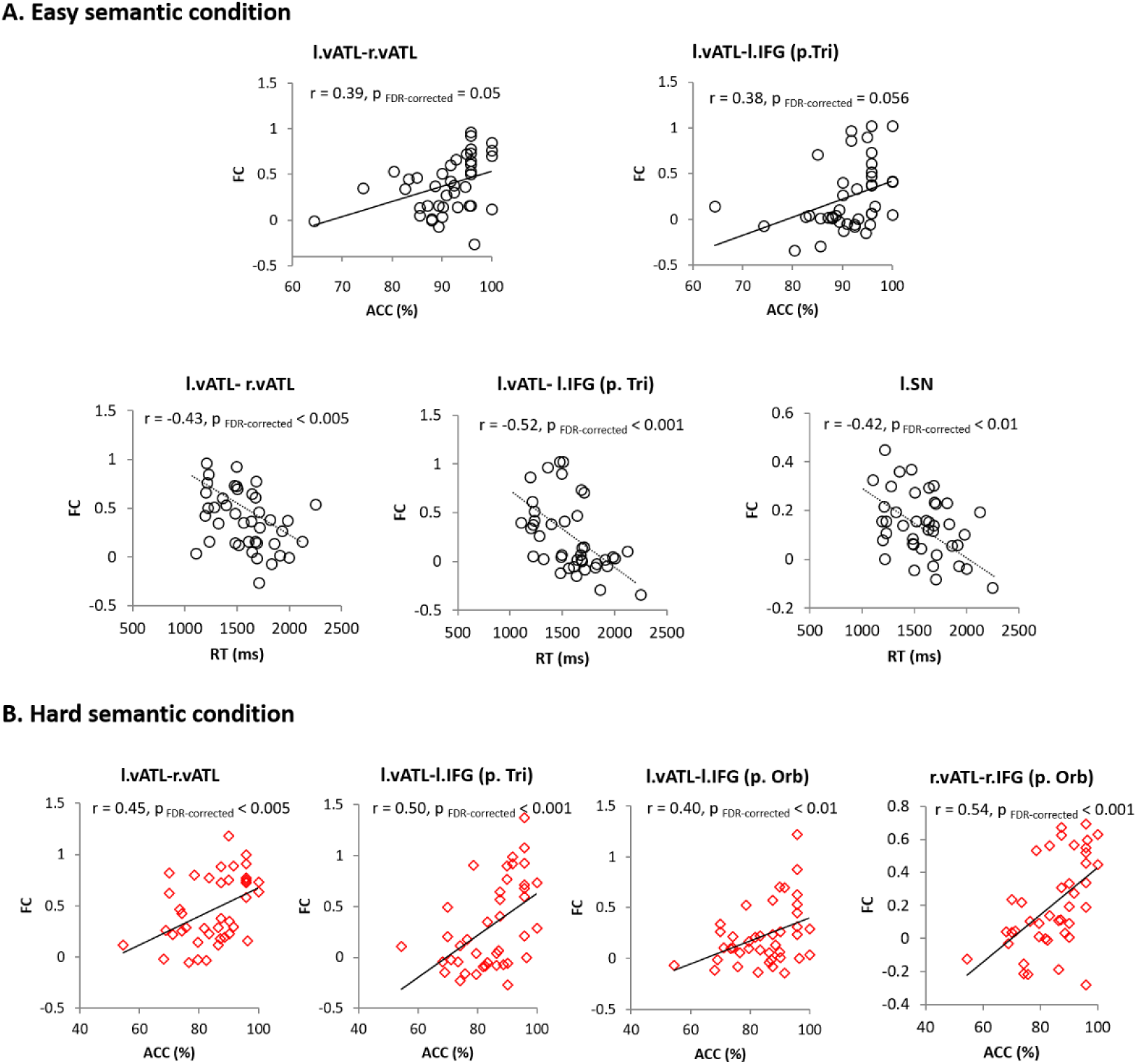
FC-behavioural correlations. A) Easy semantic condition. Black circle represent individual’s performance during the easy semantic condition. B) Hard semantic condition. Red diamond indicates each individual’s performance (accuracy) during the hard semantic condition. * p < 0.05

## Discussion

The overarching aim of this study was to begin the examination of the mechanisms that make higher cognitive systems well-engineered and thus resilience to variable performance demands, perturbation and minor damage. Beyond investigations of executive function (where demand variations are central to the experimental design), other domains of higher cognition are often examined at a single performance demand level in each study (though at different levels across studies) whilst other parameters of interest are manipulated. Yet the question of how resilience is achieved in higher cognitive systems is important both for cognitive and clinical neuroscience. In this study, we explored the intrinsic neural dynamics of semantic network in healthy participants and how the system alters to be resilient to changes in performance demand. The results directly paralleled those found in previous investigations of the semantic network after neurological damage (14, 15) or TMS perturbation (21, 22, 41) suggesting that the well-engineered neurocognitive network for semantic cognition has intrinsic mechanisms that make it resilience to changing performance demands, which are also engaged when the system is partially compromised. In summary, we found that semantically-demanding conditions upregulated the left-lateralized semantic regions including the vATL, IFG and pMTG as well as other regions in domain general networks. More specifically, there was increased activation in the bilateral ATL supporting the notion that semantic representation is inherently bilateral in nature (1, 31, 42). Functional connectivity between these regions (interhemispheric vATL connectivity and the vATL–PFC) was also increased. Importantly, this strengthened connectivity was associated with better semantic task performance. Our results suggest that semantic cognition is founded on a flexible, dynamic system making itself resilient to internal (e.g., system damage) and external changes (e.g., task demands).

Importantly, these intrinsic resilience-related mechanisms were observed in both the domain-specific representational system and the parallel executive control networks. We observed dynamic changes in the semantic representational system: semantically-demanding conditions increased the regional activity in the bilateral vATL and functional connectivity between them, which correlated with semantic performance. Note these same demand-related areas were also found in the easy condition if the statistical threshold was reduced. This fits with the notion that there is an intrinsic broader network that can support function but, to save energy, its level of activation is titrated against performance demand (which we refer to as ‘variable neuro-displacement’ (29)). This mechanism is analogous to variable displacement in combustion engines where cylinder function is down-regulated or switched off to save energy but are re-engaged when increased performance is needed (30). The bilateral nature of ATL-reliant semantic representation could, itself be crucial for its resilient, well-engineered characteristics. Past studies of healthy participants after ATL rTMS, patients with resection for temporal lobe epilepsy, and comparative neurosurgical investigations have shown that bilateral lesions are required for substantial, chronic semantic impairments (i.e., breaking the resilience of the system) and that after unilateral stimulation or resection, there are compensatory upregulations of activation in contralateral regions, peri-damage areas and increased positive transcallosal functional connectivity (18, 19, 21, 31, 43). The resilience that follows from bilateral neural systems has also been formally demonstrated and explored through implemented neurocomputational models of bilateral semantic representation. These show that a bilaterally-supported functional network is much more resilient to unilateral damage and has greater capacity for experience-dependent plasticity-related recovery (20).

As well as these dynamic changes in the domain-specific network of interest (cf. semantic representation), we also observed upregulations in the parallel executive-control networks and their connectivity to the domain-specific representations. The assistance from the executive systems, again, reflects its inherent function in the healthy system, i.e., when processing is difficult or the representations need to be adjusted then the executive systems kick in (10, 12, 44). Here, the demanding semantic condition upregulated the IFG and pMTG plus increased functional connectivity between them as well as between the vATL and IFG. Importantly, individuals with stronger vATL-IFG connectivity performed the semantic task better in both easy and hard conditions.

These types of changes and their associated explanations, directly mirror previous patient (14, 15) and TMS explorations (21, 22). This would seem to imply that the dynamic changes observed following brain damage or perturbation may not reflect *de novo* mechanisms that are triggered by brain damage but rather reflect the inherent mechanisms found in a well-engineered, resilient cognitive system. Turning to the engine analogy again, this is the same running the engine at higher demand levels (and not downregulating) when a part of the engine is compromised. After brain damage or TMS perturbation in the semantic system, the upregulation of the remaining representational and executive-control regions was observed and associated with the remaining semantic ability (21, 22, 45, 46). A recent TMS-fMRI study demonstrated the increased interhemispheric vATL-connectivity in semantic processing after perturbing the left vATL (22). Strikingly, we showed the same results in the healthy system by increasing task demand, suggesting that the same mechanism may underpin such changes in patients and TMS investigations.

These results also have potential implications for investigations of recovery in patients. Typically, patients and controls are asked to complete exactly the same task in the scanner, with the task adjusted so that patients are able to perform reasonably well within the scanner. Differences between patients and controls are assumed to reflect newly engaged regions/mechanisms and that these are the basis of recovery or compensation. The current results suggest, however, that easier versions of the same task may under-estimate the entire cognitive network and, in fact, the patients’ results might simply reflect the function of the remaining (full-engaged) cognitive network. In keeping with this hypothesis, recent patient studies have started to compare patient data against control data collected at two levels of demand (31, 47, 48). For example, Brownsett et al (48) showed that the upregulated regions observed in post-stroke aphasic patients whilst listening to clear speech aligned with regions that heathy controls upregulated when listening to degraded speech. Likewise, epilepsy patients with temporal lobe resection (either left or right) also showed similar pattern of upregulation in the intact ATL and PFC to that of healthy controls when given a shortened response time window (31). These results provide strong evidence that the compensatory functional alterations in the impaired brain might reflect intrinsic mechanisms of a well-engineered semantic system.

Finally, we note that the focus of the current study was limited to semantic system. The purpose of the current study was to explore the neural mechanism resilient to functional demands and stresses in the healthy population. Future studies will be able to assess whether these results hold across other cognitive domains, including testing the hypothesis that resilience in higher cognitive systems reflects a combination of domain-general (executive) and domain-specific networks. As well as revealing important insights about the constitution of well-engineered healthy systems, these studies should elicit potentially important insights about the neural mechanisms that support compensatory dynamic changes after brain damage or surgery.

## Funding

this research was supported by a Beacon Anne McLaren Research Fellowship (University of Nottingham) to JJ and an Advanced ERC award (GAP: 670428 - BRAIN2MIND_NEUROCOMP) and MRC programme grant (MR/R023883/1) to MALR.

