## Supplemental information for "The neural bases of resilient cognitive systems: Evidence of variable neuro-displacement in the semantic system"

### Table of contents

Supplementary Figures S1-9

Supplementary Tables S1-2

Supplementary Results

### Supplementary Figure 1

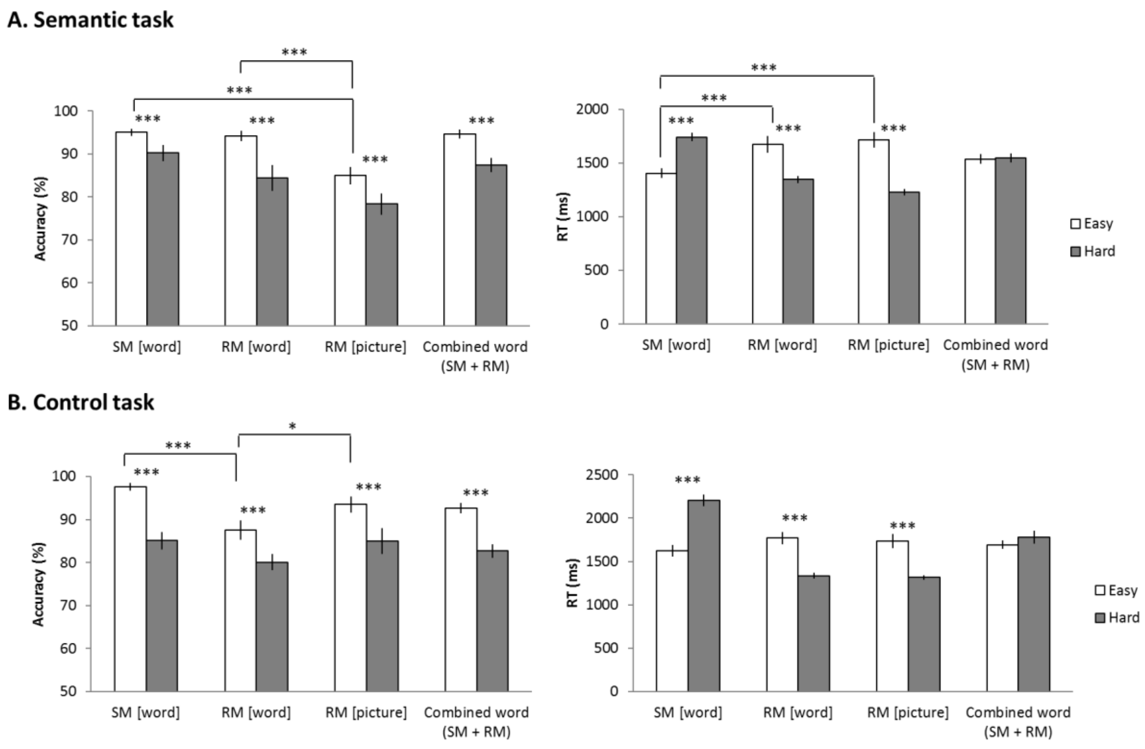


Fig. S1 Behavioural results. A) Behavioural results of semantic task. B) Behavioural results of control task. White bars represent easy condition performance and grey bras hard condition performance. Error bars indicate standard errors. ***p < 0.001, * p < 0.05.

**Supplementary Figure 2**

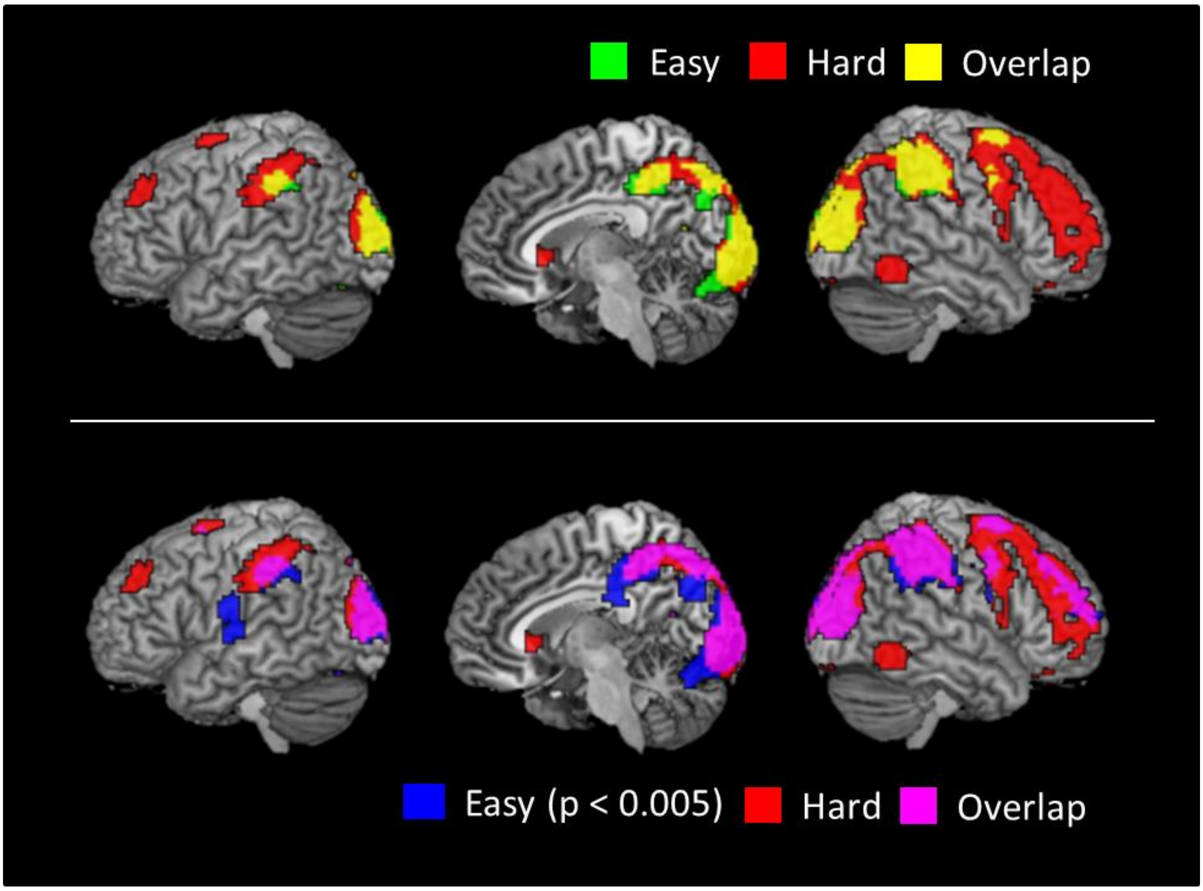


Fig. S2 fMRI results of control task (SM + RM). Red colour indicates the results of the hard semantic condition. Yellow colour represents overlapping between the easy and hard condition. Blue indicates the result of the easy semantic condition with a lower threshold ($p < 0.005$). Pink represent the overlapping between the hard condition and easy condition with a lower threshold.

**Supplementary Figure 3**

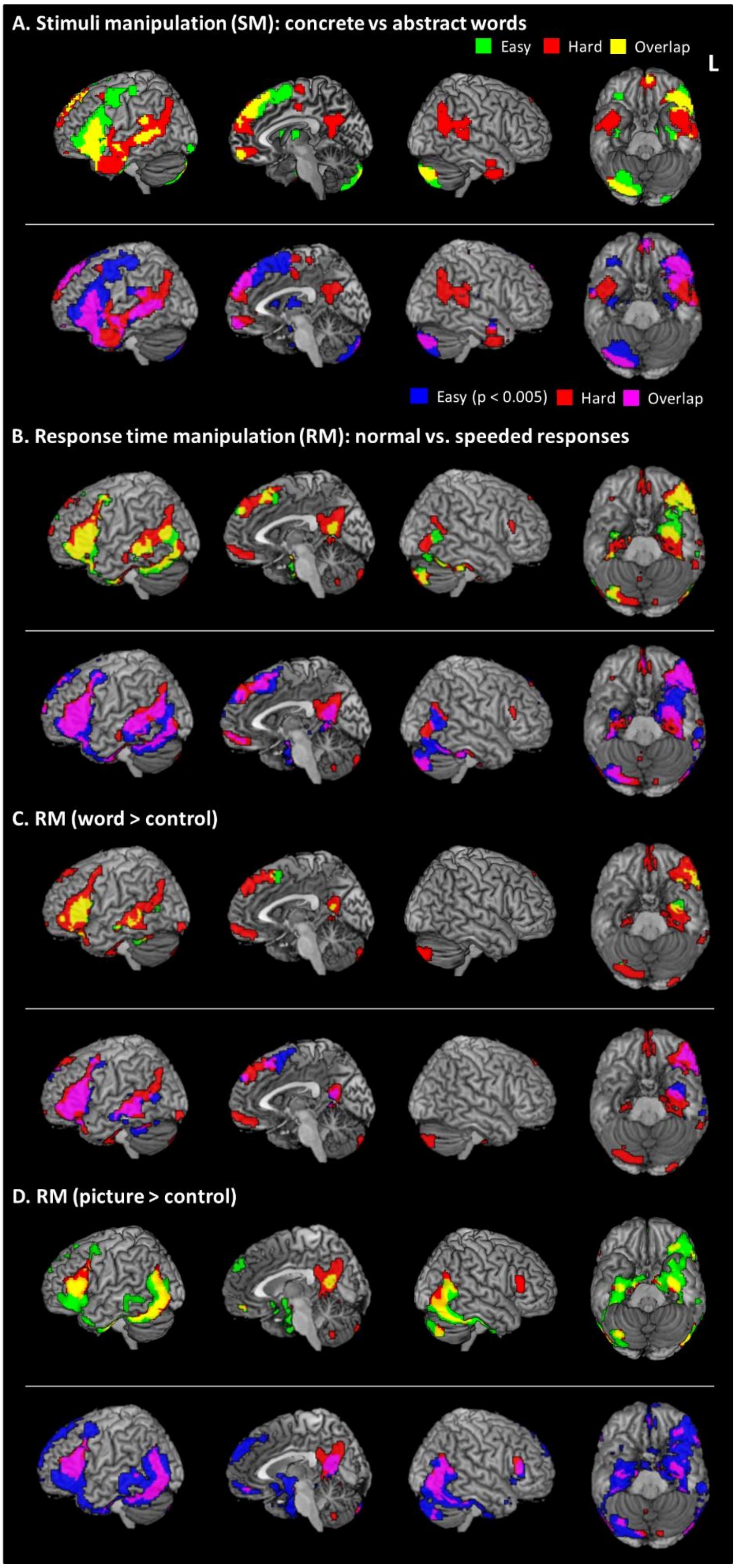


Fig. S3 fMRI results of each dataset (SM and RM) according to the modality (words and pictures) during semantic processing. A) Brain activation map of the stimulus manipulation (SM) dataset (Semantic > Control). B) Brain activation map of the response time manipulation (RM) dataset (Semantic > Control). C) Brain activation map of word modality (RM). D) Brain activation map of picture modality (RM). Green colour indicates the results of the easy semantic condition. Red colour indicates the results of the hard semantic condition. Yellow colour represents overlapping between the easy and hard condition. Blue indicates the result of the easy semantic condition with a lower threshold ($p < 0.005$). Pink represent the overlapping between the hard condition and easy condition with a lower threshold.

### Supplementary Figure 4

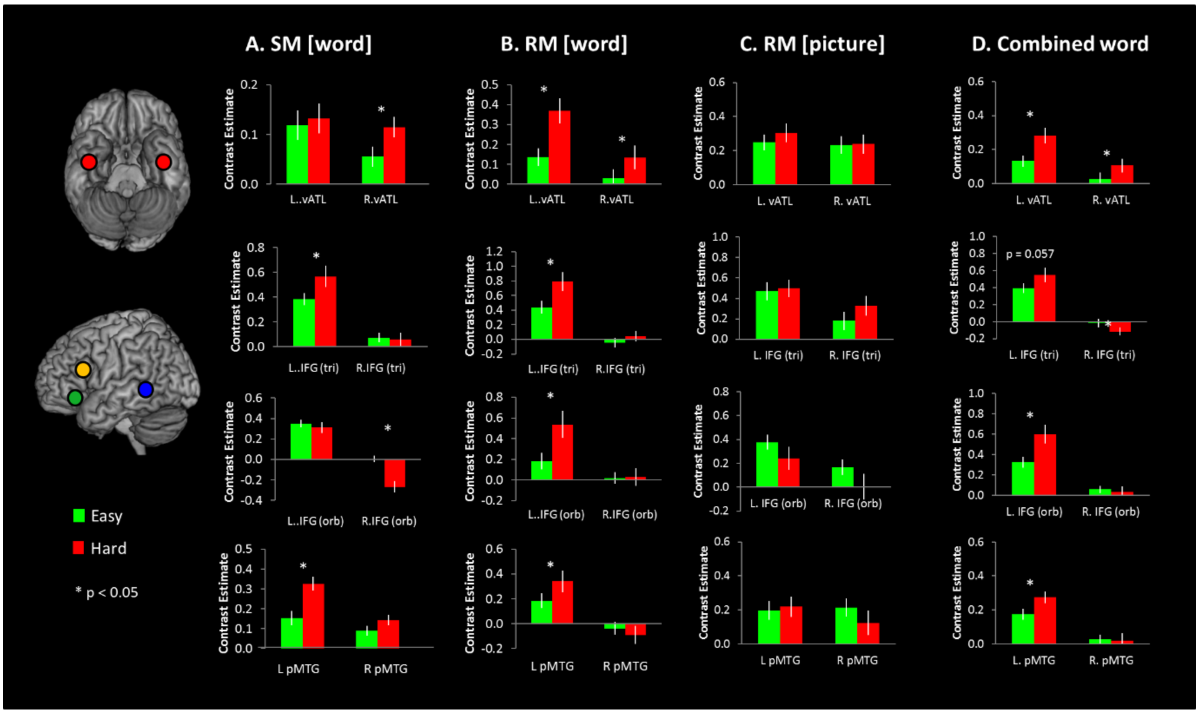


Fig. S4 ROI analysis for each dataset according to the modality. Green bars represent the easy semantic condition and red bars the hard semantic condition. * P < 0.05

**Supplementary Figure 5**

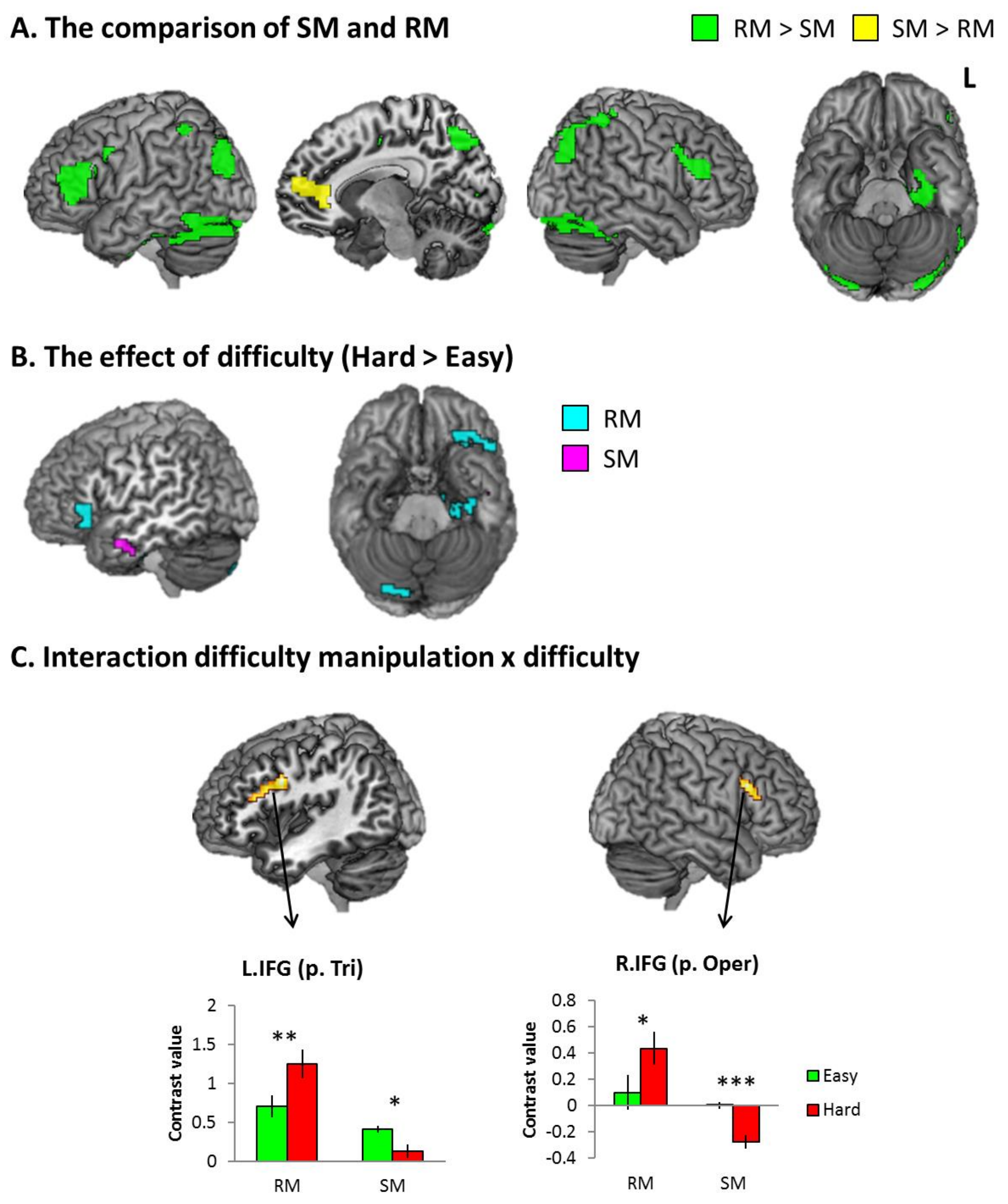


Fig. S5 The results of difficulty manipulation (SM [word] and RM [word]). A) The comparison between SM and RM. Green indicates the contrast RM > SM (semantic > control). Yellow represents the contrast SM > RM (semantic > control) B) The results of contrast (Hard > Easy) according to the manipulation. Cyan indicate the result of RM dataset. Pink represents the result of SM dataset. C) The result of interaction between the manipulation and difficulty. Green bar represents the contrast value of the easy semantic condition (semantic>control). Red bar indicates the contrast value of the hard semantic condition. Error bars indicate standard errors. *p < 0.05, ** p < 0.01, *** p < 0.001

**Supplementary Figure 6**

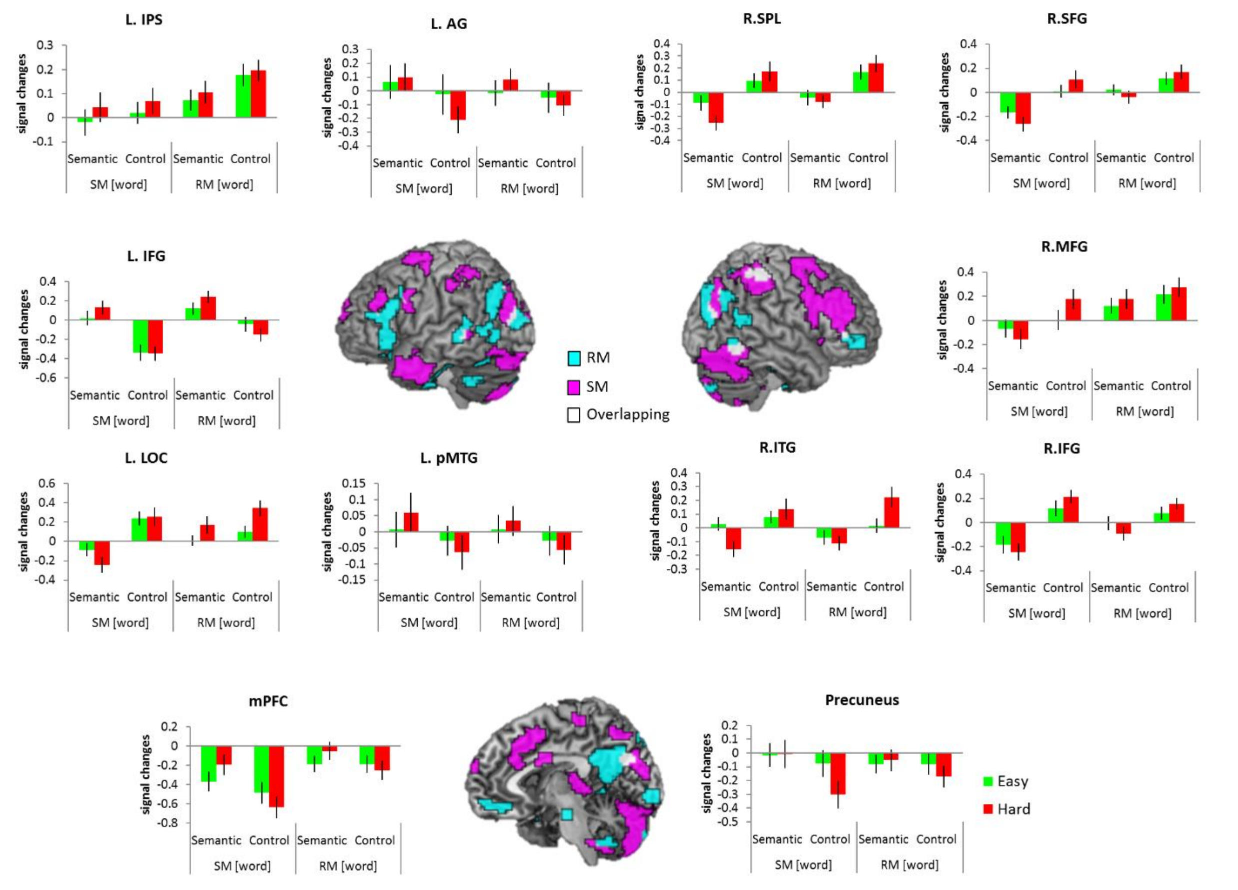


Fig. S6 Results of interaction between task and difficulty according to difficulty manipulation (SM [word] and RM [word]). Green bars represent the easy condition and red bars the hard condition. Error bars indicate standard errors.

**Supplementary Figure 7**

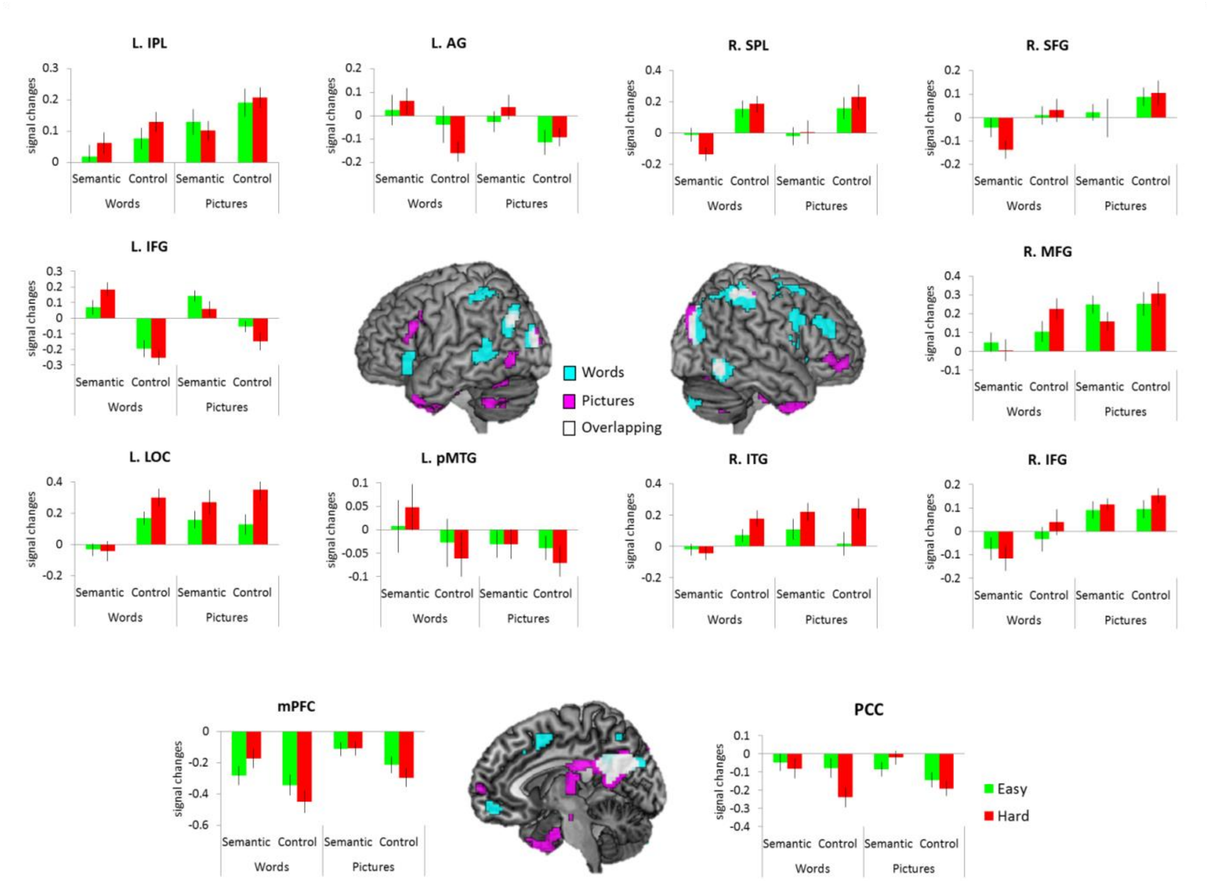


Fig. S7 Results of interaction between task and difficulty according to modality (word [SM + RM] and picture [RM]). The activation map of words was thresholded at p unc < 0.001. The activation map of pictures was thresholded at p unc < 0.01 for the display. Green bars represent the easy condition and red bars the hard condition. Error bars indicate standard errors.

### Supplementary Figure 8

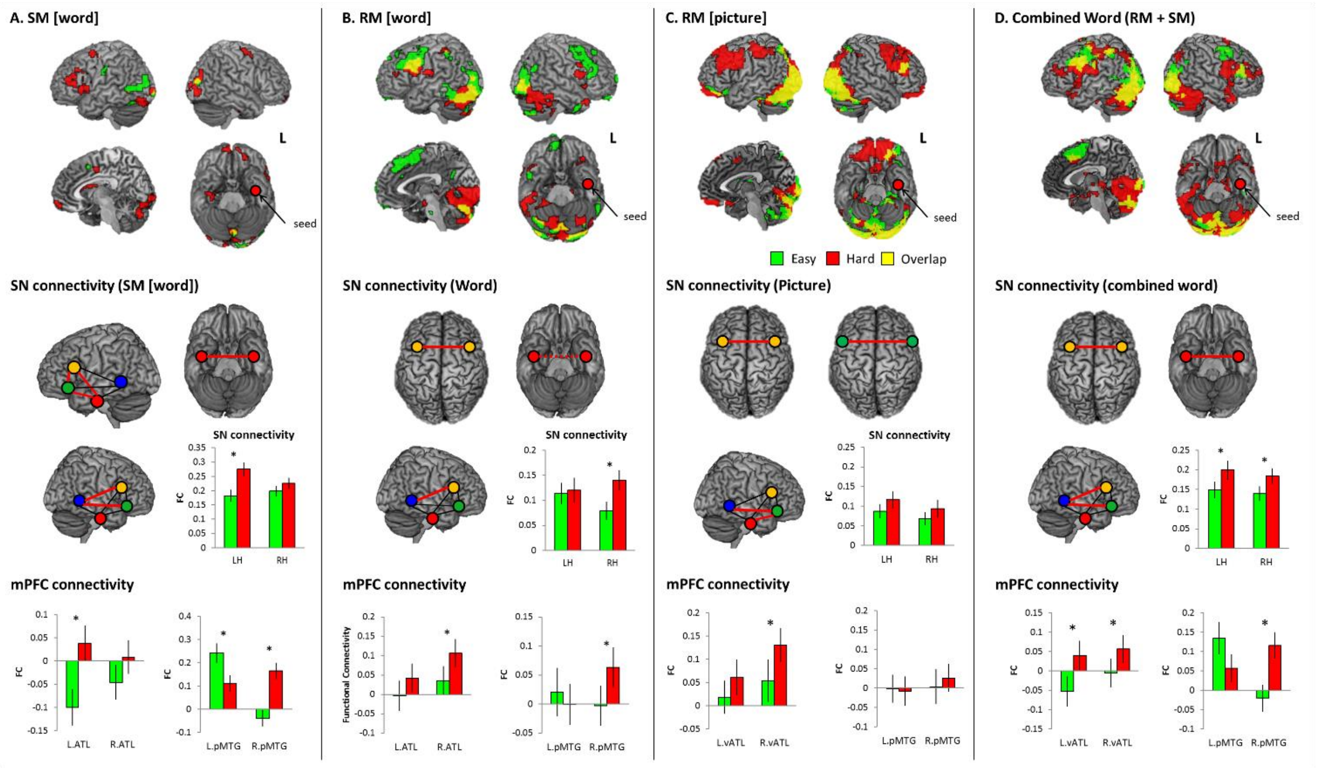


Fig. S8 Results of FC analysis. A) SM [word] results. **Top**: The results of PPI with the l.vATL seed. Green colour indicates the results of the easy semantic condition. Red colour indicates the results of the hard semantic condition. Yellow colour represents the overlapping. **Middle**: results of ROI-to-ROI analysis in the semantic network Black line represents a connection between ROIs and red line a significantly increased connection between ROIs in the hard semantic condition compared to the easy semantic condition (dotted red line, $p = 0.07$). **Bottom**: result of the mPFC connectivity. B) RM [word] results. C) RM [picture] results. D) The combined word (SM + RM) results. Note that the PPI results were displayed at p uncorrected < 0.01 at a voxel level for SM [word] and RM [word].

Error bars indicate standard errors. * $p < 0.05$

### Supplementary Figure 9

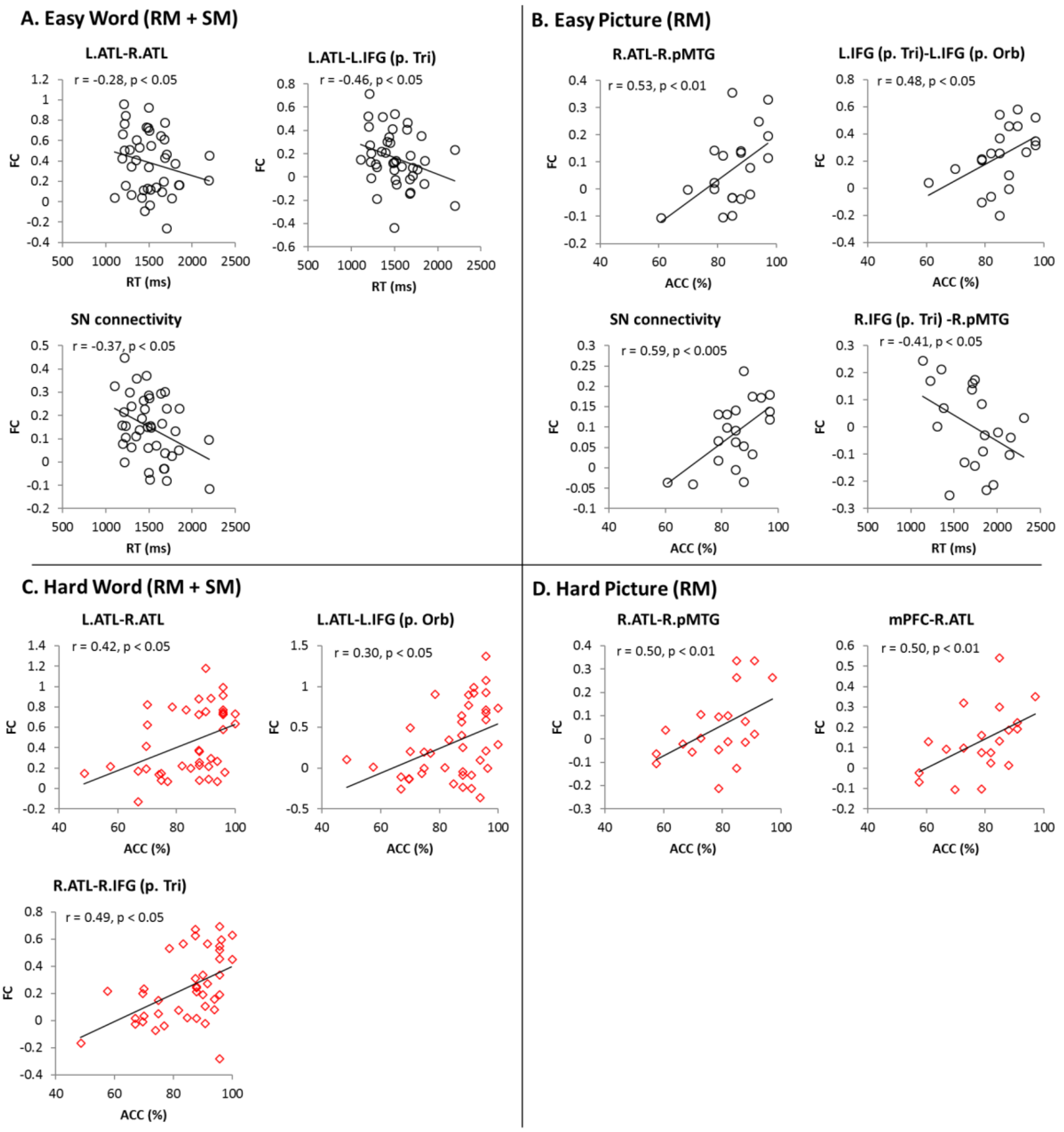


Fig. S9 FC-behavioural correlations according to modality. Black circles indicate individual semantic performance during the easy condition. Red diamonds represent individual semantic performance during the hard condition.

**Table S1**

| Contrast | Cluster size | T | x | y | z | Brain area |
|---|---|---|---|---|---|---|
| Easy Semantic > Control | 3397 | 8.16 | -57 | 27 | 6 | L IFG (p. Triangularis) |
| | | 7 | -33 | -6 | -39 | L Fusiform Gyrus |
| | | 6.88 | -51 | 33 | -3 | L IFG (p. Orbitalis) |
| | | 6.68 | -51 | 21 | -12 | L TP |
| | | 6.51 | -34 | -6 | -39 | L vATL |
| | | 6.23 | -33 | 6 | 36 | L SFG |
| | | 6.22 | -33 | 6 | 30 | L Precentral Gyrus |
| | 958 | 6.36 | -66 | -33 | -3 | L pMTG |
| | | 5.45 | -53 | -42 | 3 | L pMTG |
| | 420 | 6.55 | 39 | -78 | -36 | R Cerebelum |
| | | 6.09 | 24 | -84 | -36 | R Cerebelum |
| | 217 | 6.22 | 33 | -9 | -36 | R Fusiform Gyrus |
| | | 4.24 | 21 | -9 | -18 | R Hippocampus |
| | | 2.96 | 45 | 18 | -33 | R TP |
| | 221 | 4.91 | -3 | -54 | 9 | L Precuneus |
| | | 4.27 | -3 | -33 | 3 | L Precuneus |
| Hard Semantic > Control | 4991 | 9.24 | -57 | 24 | 3 | L IFG (p. Triangularis) |
| | | 8.51 | -51 | 27 | -3 | L IFG (p. Orbitalis) |
| | | 8.07 | -45 | 30 | -6 | L IFG (p. Orbitalis) |
| | | 7.31 | -45 | -63 | 21 | L AG |
| | | 6.59 | -39 | -15 | -30 | L vATL |
| | | 5.46 | -63 | -42 | 3 | LpMTG |
| | | 5.24 | -45 | -63 | 21 | LpMTG |
| | 546 | 7.81 | -6 | -54 | 12 | L Precuneus |
| | | 7.31 | -60 | -60 | 9 | LpMTG |
| | | 7.08 | -24 | -18 | -15 | L Hippocampus |
| | | 6.84 | -12 | 42 | 48 | L SFG |
| | | 6.79 | -42 | 27 | -21 | L TP |
| | 380 | 5.96 | 27 | -15 | -18 | R Hippocampus |
| | | 4.85 | 54 | -9 | -18 | R pMTG |
| | | 4.48 | 39 | -45 | -24 | R Fusiform Gyrus |
| | | 3.59 | 42 | -15 | -36 | R vATL |
| | | 3.27 | 45 | 21 | -33 | R TP |
| | 418 | 8.07 | 33 | -78 | -36 | R Cerebelum |
| | | 5.61 | 24 | -84 | -36 | R Cerebelum |
| | 180 | 4.91 | 57 | -66 | 24 | R Middle Occipital Gyrus |
| | | 4.57 | 60 | -60 | 18 | R pMTG |
| | | 4.52 | 69 | -39 | 3 | R pMTG |
| | | 4.37 | 63 | -57 | 9 | R pMTG |
| | | 4.3 | 54 | -66 | 39 | R AG |
| | | 3.99 | 69 | -36 | 18 | R Superior Temporal Gyrus |
| | 157 | 7.94 | 0 | 48 | -15 | L mPFC |

| | | | | | | |
|---|---|---|---|---|---|---|
| Easy Control > Semantic | 4699 | 9.88 | 27 | -72 | 21 | R Superior Occipital Gyrus |
| | | 9.12 | 33 | -75 | 15 | R Middle Occipital Gyrus |
| | | 8.18 | 24 | -63 | 33 | R Superior Occipital Gyrus |
| | 398 | 6.64 | 21 | 0 | 48 | R FEF |
| | | 4.32 | 48 | 5 | 39 | R Precental Gyrus |
| | 227 | 5.15 | -54 | -36 | 39 | L IPS |
| | | 4.97 | -35 | -42 | 39 | L IPS |
| Hard Control > Semantic | 5748 | 11.35 | 42 | -42 | 54 | R IPS |
| | | 10.29 | 30 | -78 | 27 | R Middle Occipital Gyrus |
| | | 9.88 | 24 | -66 | 39 | R Superior Occipital Gyrus |
| | | 9.67 | -36 | -42 | 39 | L IPS |
| | | 9.01 | -15 | -69 | 48 | L SPL |
| | | 8.93 | -18 | -63 | 51 | L SPL |
| | 1733 | 9.01 | 27 | -63 | 27 | R Superior Occipital Gyrus |
| | | 7.3 | 51 | 9 | 36 | R Precentral Gyrus |
| | | 7.05 | 48 | 6 | 33 | R Precentral Gyrus |
| | | 5.05 | 42 | 42 | 27 | R MFG |
| | | 5.63 | 54 | 9 | 12 | R IFG (p. Opercularis) |
| | | 5.14 | 36 | 36 | 45 | R MFG |
| | | 5.07 | 33 | 60 | 18 | R SFG |
| | 222 | 7.03 | -21 | 3 | 63 | L SFG |

Table S1 Result of whole brain analysis

**Table S2**

| Cluster size | F | x | y | z | Brain area |
|---|---|---|---|---|---|
| 731 | 56.24 | 42 | -42 | 54 | R IPS |
| | 31.91 | 30 | -81 | 33 | R LOC |
| | 25.52 | 30 | -57 | 48 | R AG |
| | 25.44 | 54 | -30 | 54 | R IPS |
| | 25.01 | 12 | -69 | 48 | R PCC |
| | 21.02 | 36 | -60 | 54 | R SPL |
| | 20.77 | 42 | -81 | 27 | R LOC |
| 509 | 40.59 | -36 | -42 | 39 | L IPS |
| | 30.15 | -24 | -60 | 51 | L SPL |
| | 28.59 | -21 | -57 | 48 | L SPL |
| | 23.28 | -30 | -51 | 33 | L AG |
| 321 | 23.4 | -3 | -69 | 30 | L Cuneus |
| | 21.21 | -9 | -78 | 30 | L Cuneus |
| | 19.14 | -3 | -48 | 30 | L PCC |
| | 13.36 | -6 | -51 | 15 | L PCC |
| 183 | 21.86 | 33 | 18 | 60 | R MFG |
| | 21.47 | 27 | -6 | 60 | R SFG |
| | 19.98 | 30 | 6 | 60 | R MFG |
| 166 | 29.69 | 54 | 12 | 36 | R Precentral Gyrus |
| | 21.94 | 45 | 3 | 30 | R Precentral Gyrus |
| | 19.85 | 60 | 18 | 30 | R IFG (p. Opercularis) |
| | 18.86 | 54 | 9 | 15 | R IFG (p. Opercularis) |
| | 18.78 | 48 | 0 | 24 | R Precentral Gyrus |
| 111 | 20.13 | -51 | -27 | 0 | L pMTG |
| | 17.03 | -57 | -36 | -3 | L pMTG |
| | 15.34 | -66 | -45 | 9 | L pMTG |
| 94 | 21.01 | -48 | -63 | 27 | L AG |
| | 20.98 | -51 | -69 | 39 | L AG |
| 92 | 25.63 | -39 | -87 | 15 | L LOC |
| 91 | 36.1 | 57 | -57 | -9 | R ITG |
| 79 | 24.19 | 45 | 39 | 27 | R MFG |
| 65 | 25.66 | 0 | 51 | -12 | L mPFC |
| 62 | 20.5 | -9 | 9 | 51 | L SMA |
| | 15.04 | 6 | 15 | 48 | R SMA |

Table S2 Results of interaction between task and difficulty

### Supplementary Results

1. Behavioural results

In order to examine the effect of modality, ANOVAs with task and difficulty were performed according to the difficulty manipulation (SM and RM) and modality (words and pictures). In accuracy, there was a significant main effect of task in the RM [word] ($F_{1, 19} = 8.68$, $p < 0.01$), RM [picture] ($F_{1, 19} = 12.40$, $p < 0.01$) and the combined word (SM +RM) ($F_{1, 40} = 6.94$, $p < 0.05$) and a main effect of difficulty in all modalities [RM [word] ($F_{1, 19} = 11.96$, $p < 0.005$), RM [picture] ($F_{1, 19} = 21.49$, $p < 0.001$), combined words ($F_{1, 40} = 37.51$, $p < 0.001$)]. *Post hoc* paired t-tests for the semantic task revealed that accuracy was significantly decreased in the hard condition for all modalities and the picture modality was more difficult than the words even in the easy condition ($p < 0.001$) (Fig. S1A). The control task showed that the hard condition had significantly lower accuracy in all modalities and the RM [word] easy condition showed the decreased accuracy compared to SM [word] and RM [picture] (Fig. S1B). In RT, there was a significant main effect of task in RM [picture] ($F_{1, 19} = 4.89$, $p < 0.05$) and the combined word ($F_{1, 40} = 22.48$, $p < 0.001$), a main effect of difficulty in the RM [word] ($F_{1, 19} = 59.69$, $p < 0.001$) and RM [picture] ($F_{1, 19} = 48.98$, $p < 0.001$). *Post hoc* paired t-tests for the semantic task revealed that the RT was significantly decreased in the RM [word] and RM [picture] hard condition due to the response time manipulation and, in easy condition, the RM conditions was slower than the SM ($p < 0.05$). In the control task, the difficulty manipulation was successful in the RT: RM conditions showed the significantly reduced RT in the hard condition compared to the easy condition, whereas SM showed the opposite pattern (Fig. S1B). There was no significant difference in the combined word because the SM and RM were collapsed.

2. fMRI results

We investigated the effect of difficulty manipulation (SM and RM) and modality (word and picture) according to task difficulty. Fig. S3-7 summarised the results. The word modality evoked significant activation in the left-lateralized semantic network in the easy condition and more widespread activation in the left semantic regions as well as the right vATL, aMTG, AG, and mPFC for the hard condition, which is similar to the main findings (Fig. S3A-C). In SM, the hard semantic condition (abstract word) upregulated the superior and lateral ATL bilaterally (Fig. S3A) whereas, in RM [word], medial ATL (Fig. S3B). In contrast to the word modality, the picture modality evoked bilateral activation in the vATL and pMTG as well as the left IFG for the easy condition and additional activation in the right IFG for the hard condition (Fig. S3D). Note that all easy conditions with lower threshold ($p$ uncorrected $< 0.005$ at a voxel) showed the regions overlapping with areas upregulated in the hard conditions.

ROI analyses also showed similar results to the main findings: the regional activity in left semantic regions (vATL, IFG [p. orb, p.tri], and pMTG) significantly increased in the hard condition and only the right vATL showed significant up-regulation in all word modality (Fig. S5). However, the picture modality demonstrated that there was no significant upregulation in semantic regions during the more demanding condition.

In order to explore the difference in difficulty manipulation, we compared SM with RM in the hard semantic condition. Semantically demanding SM (abstract word) recruited the mPFC and ACC more than RM, whereas demanding RM evoked increased activity in the left ATL, bilateral IFG, IPS, AG and fusiform gyrus relative to SM (Fig. S5A). In the direct comparison between the easy and hard condition in each dataset, the difficulty effect was found in the left lateral ATL in the SM and the left IFG and ventromedial ATL in the RM (Fig. S5B). There was a significant interaction effect in the bilateral IFG, showing that semantically demanding RM upregulated these regions, whereas hard SM down regulated them (Fig. S5C).

In the word modality, several regions showed an effect of interaction between task and difficulty, many of which overlapped with the areas exhibiting the effect of task and/or difficulty (Fig. S6-7). In SM dataset, the interaction effect was observed in the left lateral ATL, left pMTG, bilateral PFC, IPS, AG, SMA and fungiform gyrus. In RM [word] dataset, we found the significant effect in the IFG, pMTG, IPL, AG, mPFC, precuneus and right ITG. The combined word (SM + RM) showed the identical pattern of activity in the interaction effect as the main results (Fig. S7). Using the same ROIs in the main analyse, we explored the interaction effect in these regions and showed the same results as the main findings (Fig. S6). Picture modality showed a significant activation in the PCC and the right temporal pole (TP) (Fig. S7). Due to smaller number of subjects for the picture modality ($n = 20$), we applied a liberal threshold ($p$ uncorrected $< 0.01$ at a voxel) to the whole brain mapping. The results revealed activation in the bilateral IFG, TP, left AG, LOC, right SPL, ITG, and mPFC. Subsequent ROI analyses demonstrated that the hard word modality was involved in the increased activation in the IFG, AG, and pMTG in the left hemisphere as well as decreased deactivation in the mPFC, whereas the hard picture modality the increased activation in the left AG, the left LOC and right ITG and decreased deactivation in the PCC (Fig. S7). These findings indicate that picture modality seems to be processed in regions associated with visuospatial processing like the control task.

3. Functional connectivity results

The PPI analyses demonstrated that more demanding semantic processing evoked increased connectivity between the vATL and bilateral PFC and left IPS across the modalities (Fig. S8). SM [word] showed the increased FC in the interhemispheric vATL connectivity and left vATL-IFG (Fig. S8A). RM [word] also showed the increased FC between the left vATL and the bilateral PFC (Fig. S8B). The combined word modality showed increased FC between the left and right ATL (superior and ventral) (Fig. S8D), whereas the picture modality between the left vATL and mPFC (Fig. S8C). ROI-to-ROI connectivity analyses demonstrated increased FC in the interhemispheric vATL, vATL-IFG [p. orb, p.tri] in the left hemisphere and IFG-pMTG in the right hemisphere in SM [word] (Fig. S8A). Overall, the left SN was upregulated in FC during abstract word processing. In RM, we found the enhanced interhemispheric connectivity in the vATL and IFG as well as IFG-pMTG in the right hemisphere (Fig. S8B). The right SN was significantly upregulated in FC during faster semantic response. The combined word modality showed the increased interhemispheric FC in the vATL and IFG [p. tri] and IFG–pMTG connectivity in the right hemisphere for the words (Fig. 8D). The picture modality showed the increased interhemispheric FC in the IFG [p. orb, p.tri] and IFG–pMTG as well as vATL–IFG [p. orb] connectivity in the right hemisphere (Fig. 8C). The SN connectivity in both hemispheres was significantly increased for the hard word modality (Fig. 8D). In the picture modality, the SN connectivity also was increased during more demanding picture processing but did not reach the significance. Also, the connectivity between the mPFC and semantic regions (vATL and pMTG) demonstrated the increased FC in the hard semantic condition across the modality and dataset (Fig. 8A-D).

We also investigated the FC-behavioural correlations for each modality. The word modality revealed the similar results to the main findings. Individuals with stronger FC in the interhemispheric vATL (left–right vATL) connectivity and vATL–IFG connectivity in the both hemispheres performed the hard word condition better (higher accuracy) (Fig. S9A & C). The picture modality demonstrated that, in the easy condition, there was positive correlations between RT and vATL-connectivity, vATL-IFG as well as SN connectivity (Fig. 9B). In the hard condition, the FC in the vATL-connectivity, right vATL–right pMTG, right IFG- right pMTG and SN connectivity were positively correlated with the semantic performance (Fig. S9D). The results from each dataset showed similar trends but did not reach the significance.
